# Within-host evolution of *Klebsiella spp.* from intestinal carriage to bacteremia

**DOI:** 10.64898/2026.01.05.697662

**Authors:** Cécile Gosset, Olaya Rendueles, Nicolas Charbonnel, Caroline Massit, Laurence Nakusi-Ollivier, Damien Balestrino, Richard Bonnet, Pierre Saint-Sardos, Christiane Forestier, Bertrand Souweine, Sylvie Miquel

## Abstract

Gut colonization by the Gram-negative bacillus *Klebsiella pneumoniae* is a significant risk factor for extra-intestinal infections. However, the mechanisms by which this opportunistic pathogen causes disseminated infections remain poorly understood. To investigate this phenomenon, 22 strain pairs of *Klebsiella spp*. were sequentially isolated from rectal swabs and blood samples of ICU patients. All strains adhered to intestinal epithelial cells *in vitro*, but failed to invade or disrupt epithelial barrier integrity. Few phenotypic differences between strains of the same pair were observed regarding their biofilm-forming capacities and ampicillin resistance. Whole genome sequencing of five pairs showed diverse sequence types, the presence of numerous antibiotics resistance genes but few virulence genes. Pairwise comparison of genomic sequences of blood and fecal isolates evidenced a few single nucleotide polymorphisms and small insertions-deletions, mostly affecting genes involved in biosynthesis of surface structures (such as capsules, pili). Most genetic changes were driven by horizontal gene transfer events, with the notable acquisition of a plasmid that enhanced bacterial fitness by eliminating competitors. In addition, some blood isolates had reduced the number of antibiotic resistance genes, underscoring the high plasticity of the *Klebsiella* resistome. Finally, one bloodstream isolate carried a *mutS* mutation, conferring a hypermutator phenotype that could increase evolvability despite the fitness burden. Together, these findings indicate that within-host genetic adaptations, rather than the acquisition of virulence traits, can enhance the colonization, competitiveness and ultimately the ability of *K. pneumoniae* to cause invasive infections.

**Importance:** *Klebsiella* commonly colonizes the human gut, and this carriage can progress to infections, particularly in intensive care unit (ICU) patients. To better understand the *in vivo* evolution, genetically associated *Klebsiella* strains isolated from the feces and blood of ICU patients were analyzed. Although strains were able to adhere to intestinal cells, none could invade or damage the intestinal barrier. Genomic comparisons of pairs strains showed limited genetic differences, mostly mutations affecting surface structures. Frequent horizontal gene transfer events were observed, which could improve bacterial competitiveness and permits a remarkably flexible resistome. Overall, this study shows that within-host evolution, rather than the gain of virulence traits, can enhance the ability of *Klebsiella* to persist, outcompete other bacteria, and ultimately cause infections.

## Introduction

The enterobacterial *Klebsiella* species is ubiquitous in nature and present in humans as a saprophyte in the nasopharynx and in the intestinal tract (1). *Klebsiella spp.* are part of the ESKAPEE (*Enterococcus*, *Staphylococcus aureus*, *K. pneumoniae*, *Acinetobacter baumannii*, *Pseudomonas aeruginosa, Enterobacter spp.* and *Escherichia coli*) organisms, which are the leading cause of healthcare-acquired infections worldwide (2). An increasing proportion of *Klebsiella spp.* strains have acquired antibiotic resistance and infections caused by the strains are now a public health issue. For instance, more than a third of the *K. pneumoniae* isolates reported to the European Antimicrobial Resistance Surveillance Network for 2021 were resistant to at least one of the antimicrobial groups under surveillance (fluoroquinolones, third-generation cephalosporins, aminoglycosides and carbapenems) (3). In addition to the ability to gain antibiotic resistance genes, *Klebsiella spp.* can acquire different virulence factors, which facilitates the evolution from colonization to an infectious process by modulating the immune system, thereby providing a metabolic advantage to the bacteria or promoting toxin production (4).

*K. pneumoniae* colonization is a significant risk factor for infection in ICU with ∼50% of *K. pneumoniae* infections, mostly pneumonia, resulting from patients’ own microbiota (5). Additionally, the density of intestinal colonization was shown to be correlated with the risk of subsequent infection, especially bloodstream infections, in ICU and hospitalized patients (6–10). However, the mechanisms by which *Klebsiella* is able to progress from gut commensal to pathogen are still debated. The underlying hypotheses are either dissemination via the oropharyngeal sphere caused by the presence of medical devices, or direct intestinal translocation related to the modulation of intestinal barriers (mucus and epithelium) in a polymicrobial and immunocompromised context (11–13). The latter hypothesis is supported by the fact that the intestinal barrier can be altered by various conditions such as inflammatory state, cirrhosis, drugs and gut ischemia, which disrupt the tight junctions and hence facilitate bacteria translocation (11, 14–17). In addition, some pathogenic bacteria can create their own passage through the intestinal barrier. However, the possible paracellular passage of *Klebsiella* is poorly documented and published data are conflicting (18, 19).

One promising approach to understand the pathophysiological process associated with *K. pneumoniae* dissemination within the host is to characterize and compare clinical strains isolated from feces and blood samples of patients collected during the course of their pathology using recent phylogenomic studies that have provided important insight into pathogen evolution from a macroevolutionary perspective (20). More recently, at the microevolutionary level, numerous laboratory evolution experiments have shed light on the first genetic changes leading to pathogen adaptation to different environments, including those close to clinical context such as exposure to antibiotics and macrophages (21). Some of these changes were also identified in natural populations, confirming the relevance of such experimental set-ups (22). A recent study investigated the parallel between *K. pneumoniae* evolution and the modulation of its virulence factors during a hospital outbreak (23). However, it did not allow for a direct comparison of related strains involved in disseminated infection within a single host.

In this study, we isolated 22 couples of *Klebsiella spp.* strains from ICU patients who had developed *Klebsiella* bacteremia. Experimental approaches and bacterial genome evolution analyses were combined to identify changes between and within patient isolates. The ability of all isolates to adhere to and invade intestinal cells, to cross the intestinal barrier, to form biofilm and to resist ampicillin was tested. Whole-genome sequencing of five strain pairs enabled us to link genetic changes to key phenotypic traits, including biofilm formation and antibiotic resistance. Single nucleotide polymorphisms (SNPs) were rare and predominantly occurred in genes encoding surface structures. However, extensive genomic rearrangements and horizontal gene transfer, including plasmid acquisition, were frequently observed in bloodstream isolates compared to their feces counterparts, suggesting a fast and efficient adaptation. Our study highlights that bacterial adaptation in the gut and interactions with host microbiota confer new colonization capabilities and improved fitness, leading to the emergence of bloodstream infections.

## Results

### Isolation of pairs of strains from fecal and blood samples of ICU patients

Twenty-two patients with bacteriemia due to *Klebsiella* were enrolled in the study. Demographic characteristics and comorbidities of patients, their exposure to antibiotics, vasopressor support, and organ support are given in **Table 1** and **Fig 1**. Details of the antibiotics received by these patients in the two weeks before bacteremia are shown in **Fig 1C**. The median age of the patients was 66 years (IQR 61-69) and their median SAPS II score at ICU admission was 41 (IQR 29-48). Sixteen out of the 22 patients were admitted for medical reasons, 6 others for surgical reasons, and 10/22 were treated with corticosteroids. In addition, 21/22 patients had been exposed to antibiotics within the 2 weeks prior to bacteremia. In the two weeks prior to bacteremia, 14/22 were on mechanical ventilation, 13/22 on vasopressors and 11/22 on renal replacement therapy.

**Figure 1.**
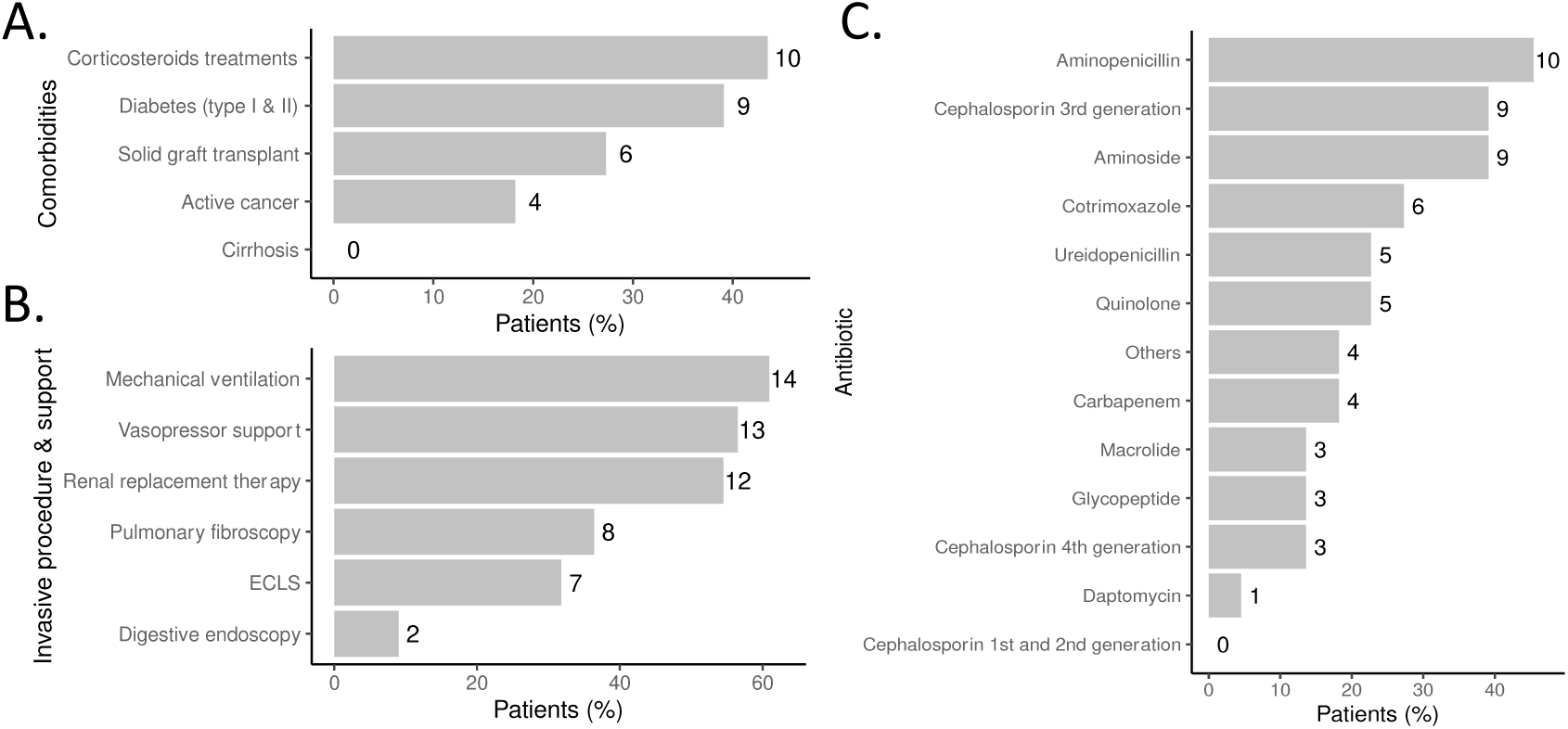
Clinical characteristics and management of intensive care unit patients with bacteremia caused by a *Klebsiella spp* strain similar to that present in their digestive carriage. **(A.)** Comorbidities (**B.)** Invasive support and (**C.)** Antibiotic treatments within 2 weeks prior to bacteriemia. ECLS: Extra-Corporeal Life Support.

**Table 1.** Demographic characteristics, comorbidities, antibiotic exposure, invasive support of intensive care unit patients identified with bacteremia caused by a *Klebsiella spp* strain similar to the one found in their digestive carriage (January 2017 and March 2021).

|  |  |
| --- | --- |
| **<u>Demographics and comorbidities</u>** |  |
| Age (years) at ICU admission (median, IQR) | 66 (61-69) |
| Male (%) | 12 (54.5) |
| BMI (kg/m<sup>2</sup>) at ICU admission (IQR) | 26 (23-30) |
| Admission for medical reasons (%) | 16 (72.7) |
| Admission for surgical reasons (%) | 6 (27.2) |
| SAPS II at ICU admission (median, IQR) | 41 (29-48) |
| **<u>Invasive procedure and support<sup>a</sup></u>** | **N = 20 (86.9%)** |
| **<u>Antibiotic exposure<sup>a</sup></u>** | **21 (95.5%)** |

ICU: Intensive Care Unit; IQR: InterQuartile Range; BMI: Body Mass Index; SAPS II: Simplified Acute Physiology Score II. <sup>a</sup> within two weeks prior to bacteremia.

A total of 85 *Klebsiella spp* isolates harboring multidrug resistance were initially collected from samples of the 22 enrolled patients. MALDI-TOF spectrometry and ERIC-PCR amplification profile analysis (**Fig S1**) identified 22 pairs of concordant strains, including 5 pairs of *K. oxytoca* and 17 of *K. pneumoniae* (**Table 2**). The overall strain antibiotic resistance profiles were recovered from patient records (**Table S1**). The Minimal Inhibitory Concentrations (MIC) of ampicillin varied from 0.2 to 125 g/L (**Table 2**). The MIC for three strains isolated from blood samples was higher than those of their feces counterpart (#3, #4 and #5) whereas the MIC for three other blood sample isolates was lower than those determined with their feces counterpart (#7, #8 and #22).

**Table 2.** Concordant strains-pairs isolated from ICU patients.

| Patients N° | Strain (stool carriage) | Ampi MIC (g/L) | Strain (blood culture) | Ampi MIC (g/L) | Species | Year of isolation |
| --- | --- | --- | --- | --- | --- | --- |
| #1 | CH1662* | 12.5+/-0 | CH1661* | 12.5+/-0 | *pneumoniae* | 2020 |
| #2 | CH1664* | 6.25+/-0 | CH1663* | 6.25+/-0 | *pneumoniae* | 2020 |
| #3 | CH1666* | 6.25+/-0 | CH1665* | 12.5+/-0 | *pneumoniae* | 2020 |
| #4** | CH1667* | 0.13+/-0,06 | CH1668* | 125+/-0 | *pneumoniae* | 2020 |
| #5** | CH1670* | 0.2+/-0 | CH1669* | 5.2+/-1.8 | *pneumoniae* | 2020 |
| #7** | CH1676 | 25+/-0 | CH1675 | 5.2+/-1.8 | *pneumoniae* | 2020 |
| #8** | CH1679 | 40.6+/-10.8 | CH1678 | 4.7+/-1.8 | *oxytoca* | 2020 |
| #9 | CH1681 | 9.4+/-4.4 | CH1682 | 9.4+/-4.4 | *pneumoniae* | 2020 |
| #10 | CH1684 | 18.8+/-8.8 | CH1685 | 18.8+/-8.9 | *pneumoniae* | 2019 |
| #11 | CH1688 | 9.4+/-3.6 | CH1687 | 9.4+/-3.6 | *pneumoniae* | 2019 |
| #12 | CH1689 | 12.5+/-0 | CH1690 | 12,5+/-0 | *oxytoca* | 2019 |
| #13 | CH1691 | 18.6+/-9 | CH1692 | 18.6+/-9 | *pneumoniae* | 2019 |
| #14 | CH1694 | 18.8+/-8.9 | CH1695 | 18.8+/-8.8 | *pneumoniae* | 2019 |
| #15 | CH1700 | 12.5+/-0 | CH1701 | 16.7+/-7.2 | *pneumoniae* | 2018 |
| #16 | CH1703 | 9.4+/-4.4 | CH1704 | 9.4+/-4.4 | *pneumoniae* | 2018 |
| #17 | CH1707 | 12.5+/-10.8 | CH1708 | 10.4+/-3.6 | *pneumoniae* | 2017 |
| #18 | CH1710 | 12.5+/-0 | CH1713 | 12.5+/-0 | *pneumoniae* | 2017 |
| #19 | CH1715 | 12.5+/-0 | CH1716 | 16.7+/-7.2 | *oxytoca* | 2017 |
| #20 | CH1720 | 20.8+/-7.2 | CH1723 | 25+/-0 | *pneumoniae* | 2017 |
| #22 | CH1724 | 25+/-0 | CH1725 | 12.5+/-0 | *oxytoca* | 2017 |
| #25 | CH1728 | 4.7+/-2.2 | CH1729 | 6.2+/-0 | *pneumoniae* | 2016 |
| #26 | CH1731 | 7.8+/-6.6 | CH1730 | 7.8+/-6.6 | *oxytoca* | 2021 |

MIC: Minimal Inhibitory concentration of Ampicillin in g/L. \*\*Strain pairs with more than one variation dilution for MIC with Ampicillin; \*Sequenced strains.

### Biofilm-forming *Klebsiella spp.* adhere to intestinal epithelial cells without inducing invasion and epithelial barrier destabilization

All strains isolated from rectal swabbing were able to adhere to intestinal Caco-2 cells at the same level as the reference AIEC LF82 strain (**Fig 2A**). After 6h of co-incubation of bacteria and Caco-2 cells, the TEER of the cell monolayer was not significantly affected by the presence of bacteria (*Klebsiella* or a commensal *E. coli* strain MG1655) (**Fig 2B**). To test whether the bacteria could be internalized by Caco-2 cells, aminoglycoside protection assays were performed, all isolates being susceptible to this antibiotic. The recovery rate of residual bacteria after amikacin treatment ranged from (95 to 2.62 10^5^ UFC/mL (**Fig 2C**). For most *Klebsiella* strains, this rate was comparable to the number of CFU observed when using the reference AIEC LF82 strain (7.33 10^5^ UFC/mL) (except strains from patient**s** #7, #16, #19, #22, and #26). However, in the absence of cells (**Fig 2C**), amikacin treatment also gave rise to numbers of CFU close to those observed in the presence of cells, indicating that Klebsiella cells were not internalized by Caco2 and that the observed protection was probably related to low diffusion of the antibiotic in bacterial aggregates formed on the cell surface.

**Figure 2.**
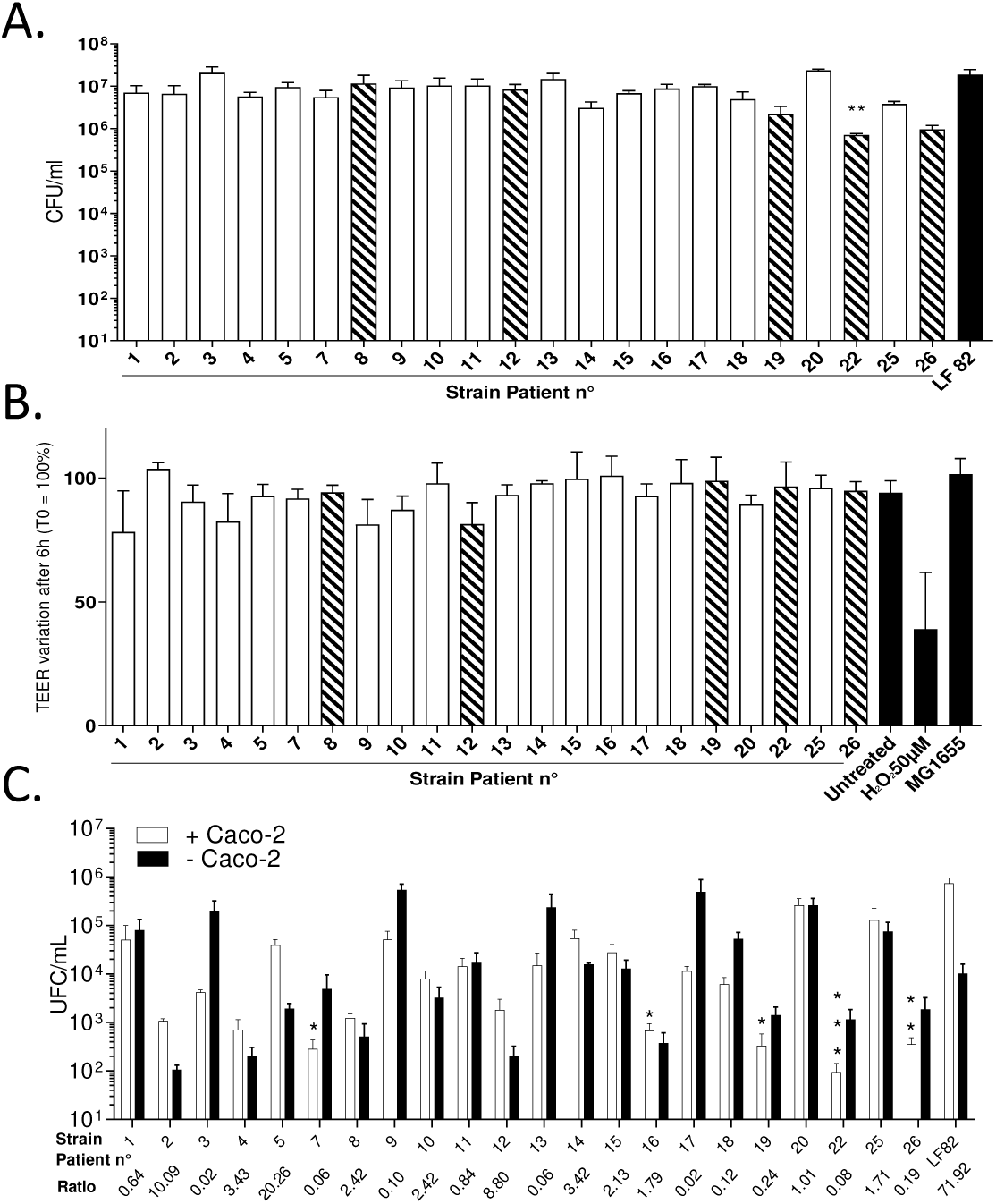
Ability of *Klebsiella* strains isolated from patient rectal carriage to interact with intestinal epithelial cells. (**A.)** Caco-2 cell-associated bacteria were quantified after centrifugation and a 4-h infection period. (**B.)** TEER measurement of a differentiated Caco-2 monolayer after 6h and 24h of bacteria and cell co-incubation. white bars: *K. pneumoniae* strains, hatched bars: *K. oxytoca* strains, black bars: controls (adherent invasive *E. coli* LF82 strain, commensal *E. coli* MG1655 strain, 50µM of H_2_O_2_ treated cells and untreated cells). (**C.)** Amikacin-based invasion assays were performed in presence of Caco-2 cells (white bars, + Caco-2) and in absence of Caco-2 cells (black bars, - Caco-2). Statistical analysis: nonparametric Kruskal-Wallis with Dunn’s multiple comparison test compared to LF82 infection condition: *p < 0.05, **p < 0.01. The invasive ability of each strain was assessed by determination of the ratio (+ Caco-2 CFU/mL / - Caco-2 CFU/mL). Each value is the mean ± SEM of at least three separate experiments. Statistical analysis: nonparametric Kruskall-Wallis with Dunn’s multiple comparison test: *P < 0.05.

### Genomic analyses of stool and bloodstream isolates show the presence of numerous ARGs but few virulence factors encoding genes

Whole genome sequencing was performed on ten isolates from patients #1 to #5 and the chronology between antibiotic treatment of the patients. Treatment details, identification of the first positive rectal carriage and the first positive blood culture are specified in **Fig 3A**. The isolates from each patient belonged to diverse sequence types, ST323, ST20, ST723, ST449 and ST45 (**Table S2**), and were phylogenetically related indicating common origins (**Fig 3B**). Each pair coded for different K and O locus types, but each strain from the same patient had the same one. None of these isolates harbored the *rmp* locus typically associated with hypervirulence phenotype (24), and they all had a low virulence score owing to the low frequency of virulence factors such as colibactin or siderophore-associated genes. Only yersiniabactin-encoding genes were identified in the isolates of 4 out of the 5 patients (**Table S2**). All strains had complete type 6 secretion system encoding genes (**Table S2**).

**Figure 3.**
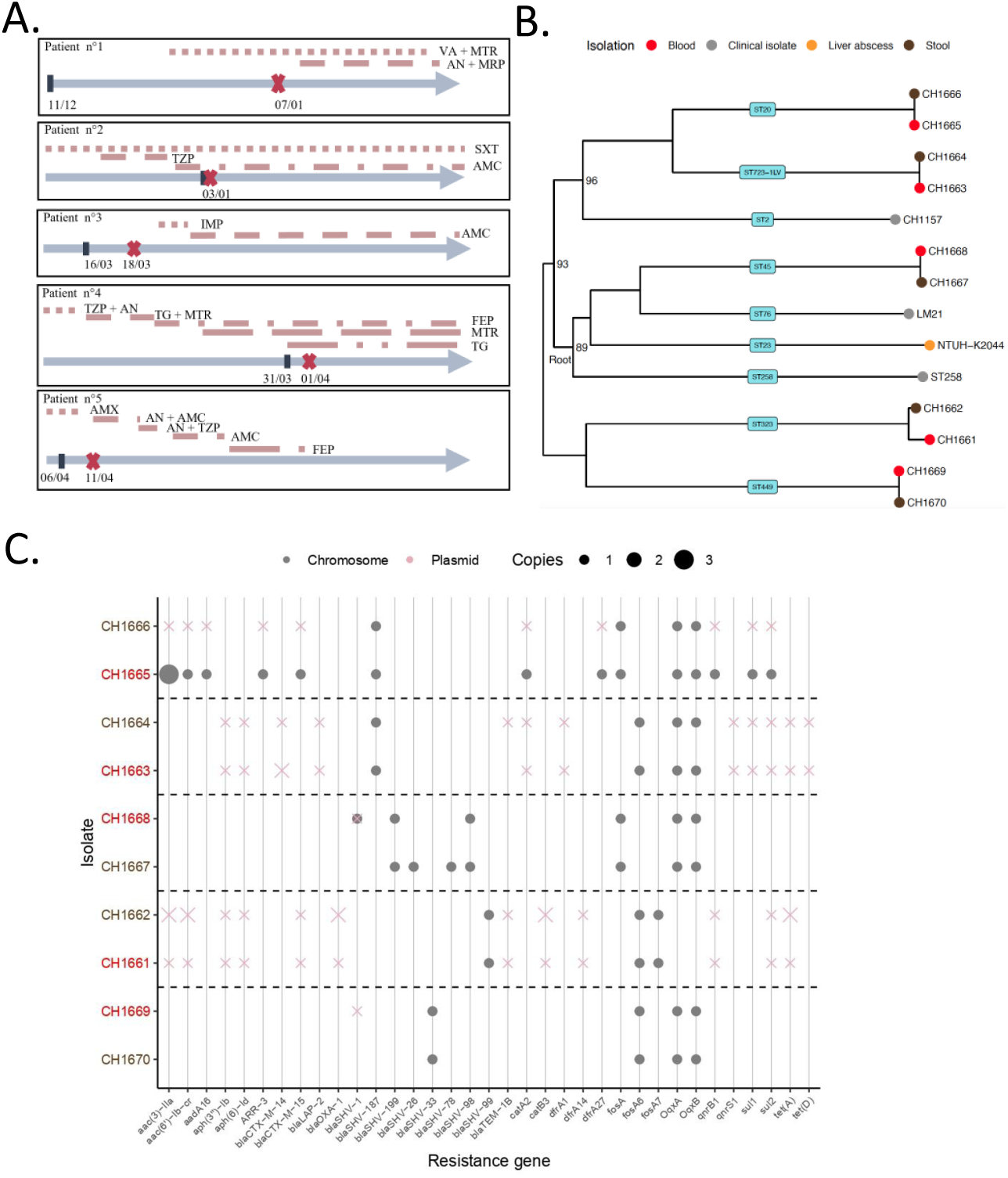
Sequence analyses of isolates from five patients. (**A.)** Timeline of patient’s antibiotic treatments and *Klebsiella* strain isolation. Intravenous antibiotic treatments are shown at the top for the five patients. Sample collection days are indicated: rectal swab (brown bars) and blood (red cross). **(B.)** Phylogenetic tree of the blood (red) and stool (brown) samples. Hypervirulent strain NTUH-K2044, multi-drug resistant NJ ST258-1, alongside two other strains previously isolated and characterized in the same hospital, strain LM21 and CH1157, were included as references. Only bootstrap values lower than 100 are indicated. **(C.)** Presence of resistance genes, as determined by ResFinder, in the different blood (red) and stool (brown) isolates. Dashed horizontal lines are used for visualization purposes to separate patients. The size of the gray dots (chromosome) and pink crosses (plasmids) indicate the number of copies.

A total of 32 extended-spectrum beta-lactamases (*bla)* genes were identified among the antibiotic-resistant genes harbored by the isolates (**Table S3, Fig 3C**), with each isolate harboring at least one copy. Half of the *bla* genes were detected within the chromosome sequences. Genomic sequences of the isolates from patient #1 included at least one copy of the following *bla* gene families, CTX, OXA, SHV and TEM. Other isolates had multiple copies of antibiotic-encoding gene (*blaOXA-1* present twice in the plasmid of strain CH1662, as were other genes conferring resistance to tetracycline -*tet*(A)- and aminoglycosides -*aac*(3)-). All 10 isolates harbored genes involved in the resistance towards quinolones, 8 coded either *oxqA* or *oxqB*, and 6 carried genes conferring resistance to ciprofloxacin (*qnrB1, qnrS1*). The number of ARGs genes per isolate did not correlate with the MIC levels of ampicillin (Spearman’s rho= 0.4, p = 0.23).

### Comparison of whole genome sequences from blood and fecal isolates detected few chromosomal changes

To investigate how within-host adaptation could shape infection, we compared whole genome sequences of the samples isolated from blood with those obtained from fecal samples of the same patients. Overall, this paired approach showed that most blood samples had few single nucleotide polymorphisms or small insertions/deletions compared to their stool counterparts (fewer than 10) (**Table S4**), with the exception of patient #1. A total of 323 SNPs were detected in the genome sequence of CH1661 strain from patient #1, compared to the sequence of his stool isolate (CH1662) (**Table S4**). These numerous mutations were correlated with the presence of a non-synonymous mutation in *mutS,* a gene involved in DNA mismatch repair, which led to a hypermutator phenotype. Mutations in genes coding for capsule production (*wzc)*, efflux pumps (*acrB*), and resistance to antimicrobials (*fusC*, *bla*) were also detected within the genome of this isolate.

The blood isolate from patient #2 (CH1663) had five chromosomal mutations compared to the genome of its counterpart from rectal swabbing (CH1664). All were in intergenic regions, except one nucleotide change, resulting in non-synonymous mutation in *robA* (encoding for an activator of the well-characterized AcrAB MDR efflux pump). Isolate from the blood sample of patient #3 (CH1665) had a premature stop in *rbsA* gene, which encodes the ribose ABC transporter ATP-binding protein. The single mutation detected in the genome of strain CH1668 from blood sample of patient #4 resulted in a frameshift within *pqqB*, a gene involved in the biogenesis of pyrroloquinoline quinone. Isolate from the blood sample of patient #5 (CH1669) had an in-frame insertion in *cpxA* gene, which encodes the sensor kinase of the CpxRA system, and a fimbriae repressor (**Table S4)**. Isolate from blood sample of patient #5 also showed a small deletion just upstream of *lysR* gene, which encodes an important virulence regulator.

Comparison of sequences of bloodstream and stool isolates also showed significant genome rearrangements. Most notably, the strain from the blood sample of patient #5 (CH1669) compared to the fecal counterpart strain, had a reversion of the *fim* switch, which controls the ON/OFF expression of type I fimbriae **(Table S4)**. This, in addition to the mutation in the fimbriae repressor *cpxA,* could explain the greater biofilm formation ability of the strain than that from fecal sample.

### Extensive horizontal gene transfer events observed within patients

We then analyzed the mobilome of the different patient isolates. Blood isolates of patients #4 and #5 had acquired novel genetic elements, namely plasmids. This represents a net gain of genes of 275 and 370 genes for strain CH1668 (patient #4) and CH1669 (patient #5), respectively, compared with the sequence of their fecal counterparts (**Table 3**). The genome from fecal isolates of patients #4 and #5 contained genes encoding for CRISPR systems (**Table 3**). No increase in the number of CRISPR arrays was observed in the sequences of their blood counterparts. The sequence of the blood isolate (CH1668) of paired strains from patient #4 had an 8.8kb plasmid (**Fig 4A**), which was absent from the stool isolate. This plasmid had 98% identity with the pSEM plasmid from *Salmonella enterica* serovar Typhimurium (25) (**Fig S5A**). A larger search for this plasmid showed that this sequence was found in over 3,000 genomes of a broad range of Enterobacteria, including *Yersinia pestis, Enterobacter cloacae* and *E. coli* (**Fig 4B**). The sequence was found in plasmids with varying sizes ranging from 5.1 kb up to 688 kb in *K. pneumoniae*, 490 kb in *K. michiganenesis*, and 355 kb in *Citrobacter freundii* (25). In our study, this 8.8 kb sequence was found in patient #4 (CH1668) as a plasmid but also in the chromosome in the form of an integrative and conjugative element (ICE) of ⁓160 kb, which codes for all the mobility machinery, including *tra* genes (**Fig S5B**). This sequence had an overall identity of 88% with one of the two plasmids acquired by strain CH1669 from patient #5. This plasmid of ⁓170 kb, which also has a phage in it, not only coded for the *bla-SHV-1* but also for the whole *fec* operon involved in the iron metabolism and operons associated with heavy metal resistance, including copper, arsenite and silver (**Fig 4C**). The backbone of the ICE was also found in the plasmids identified in strains from all other patients (**Fig S5B**). Strain CH1669 (patient #5) also gained a small plasmid of 6.6 kb (CH1669 plasmid # 2) that carried genes with unknown function (**Fig 4D**). Search of NCBI detected only one plasmid with high identity and over 50% coverage, the p6 plasmid of *Klebsiella grimontii* KOX 60 (CP067439.1) isolated in a hospital effluent in Brazil. Unlike the CH1669 plasmid #2, p6 harbors several resistance genes against aminoglycosides (26). The analysis of the mobilome of these strains underscores the ability of *K. pneumoniae* to readily exchange genetic information with other gut microbiome species, which can in turn result in competitive advantages or increased tolerance to stress.

**Figure 4.**
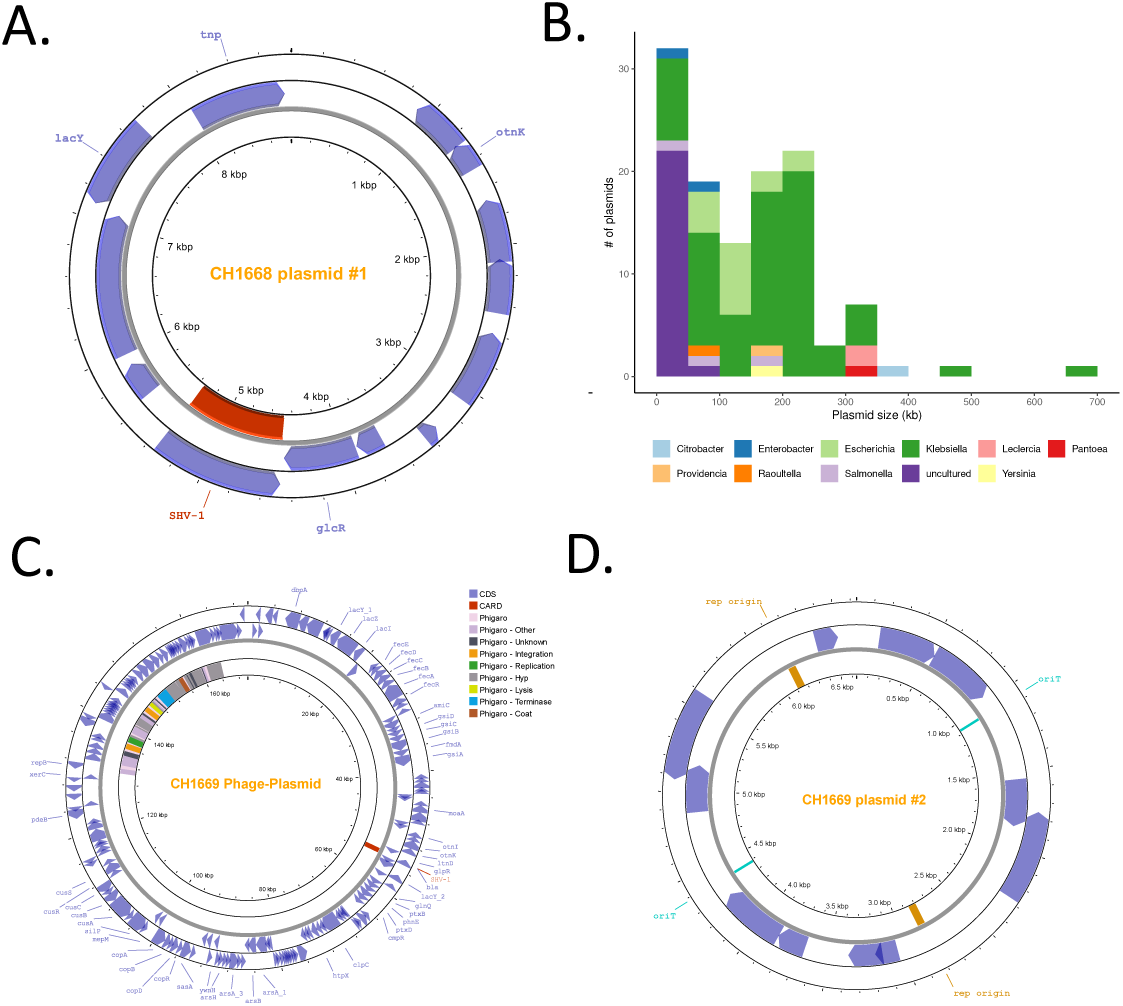
Annotation of plasmids and visualization of strains isolated from blood sample of patients #4 and #5. **(A.)** Plasmid annotation and visualization of strain CH1668. **(B.)** Plasmids with high identity to the CH1668 plasmid were also found in other plasmids across different genera of Enterobacteria, and in plasmids varying in size. **(C.)** Plasmid # 2 of strain CH1669 containing a phage. **(D.)** Plasmid # 2 of strain CH1669. Plasmids were annotated with Bakta, and visualized with Prokasee. Additionally, Phigaroo, CARD and pLAnnotate, included in Prokasee, were used to indicate plasmid features of interest.

**Table 3.** Summary statistics of the sequenced isolates.

| Patient | Isolate | Genome length (pb) | GC (%) | # contigs | CDS | Protein families | CRISPR arrays | Plasmid maps |
| --- | --- | --- | --- | --- | --- | --- | --- | --- |
| #1 | **B: CH1661** | 5510746 | 57.3 | 3 | 5099 | 4956 | 0 | **Figure S2** |
|  | **RS: CH1662** | 5522413 | 57.3 | 3 | 5237 | 5126 | 0 | Not provided |
| #2 | **B: CH1663** | 5431946 | 57.3 | 2 | 5128 | 5003 | 0 | **Figure S3** |
|  | **RS: CH1664** | 5431091 | 57.3 | 2 | 5128 | 4974 | 0 | Not provided |
| #3 | **B: CH1665** | 5556730 | 57.3 | 2 | 5128 | 5002 | 0 | **Figure S4** |
|  | **RS: CH1666** | 5547402 | 57.3 | 3 | 5254 | 5112 | 0 | Not provided |
| #4 | **B: CH1668** | 5437554 | 57.4 | 2 | 5291 | 5131 | 1 | **Figure 4** |
|  | **RS: CH1667** | 5259161 | 57.5 | 1 | 4921 | 4793 | 1 | Not provided |
| #5 | **B: CH1669** | 5457093 | 57.4 | 3 | 5218 | 5059 | 2 | **Figure 5** |
|  | **RS: CH1670** | 5280358 | 57.6 | 1 | 4943 | 4808 | 2 | Not provided |

B: Blood, RS: Rectal swab. Protein families were determined using a threshold of 99% of identity. Reducing the percentage identity to either 95% or 80% does not alter the results qualitatively. Plasmids were annotated with Bakta (63), and visualized with Prokasee (64). Additionally, Phigaroo, CARD and pLAnnotate included in Prokasee were also used to identify plasmid features of interest. Statistics provided by annotation pipeline prokka with default settings

### Within-patient evolution shows a highly dynamic resistome

The genetic analyses showed a remarkable dynamism of antimicrobial resistance genes. A plasmid rearrangement resulting in the loss of several ARGs, including a copy of *bla*OXA-1, was observed in the genome from the blood isolate of patient #1 (CH1661) compared to its fecal counterpart (**Fig S2C**) but with no concomitant reduction in the MIC of ampicillin. The plasmid from the blood strain of patient #2 (CH1663) carried an extra copy of the *bla*CTX-M-14 (**Fig 3C**) gene compared to the genomic sequence of its counterpart fecal isolate. Comparison of the genomic sequences of isolates from patient #3 indicated that one of the two plasmids present in the fecal isolate genome had integrated the chromosome of the blood isolate (**Table 3, Fig S4**). We observed a high dynamism of antibiotic resistance genes across isolates from the same patients. As expected, there was a gain of certain resistance genes. For example, in the sequence of the CH1668 blood isolate of patient #4, *blaSHV−1* was acquired both in a plasmid and in the chromosome (**Fig 3C**), which could explain the large increase in ampicillin MIC of the blood sample (**Table 2**). However, this observation cannot be extended to other samples. Other than the acquisition of antibiotic resistance genes, we also observed ARG exchanges between the chromosome and the episomes. For example, there was a chromosomal integration of a large region coding for several ARGs (patient #3). In all but one patient (#3), we also observed the absence of antibiotic resistance genes in bloodstream isolates compared to their respective stool isolate**s**. This is particularly interesting because among the five patients whose samples were sequenced, patient #3 was the only one not to receive any antibiotic treatment prior to the onset of bacteremia.

### Blood culture isolate shows efficient biofilm formation and competitivity

We then compared how the blood samples had phenotypically evolved from those isolated from the stool. All 22 strain pairs were able to form biofilm on abiotic surface**s** (**Fig 5A**). Twelve strains isolated in blood samples formed larger biofilm than their counterpart isolates from rectal swabs with a significant difference for strain pairs isolated from patients #5, #11 and #19.

**Figure 5.**
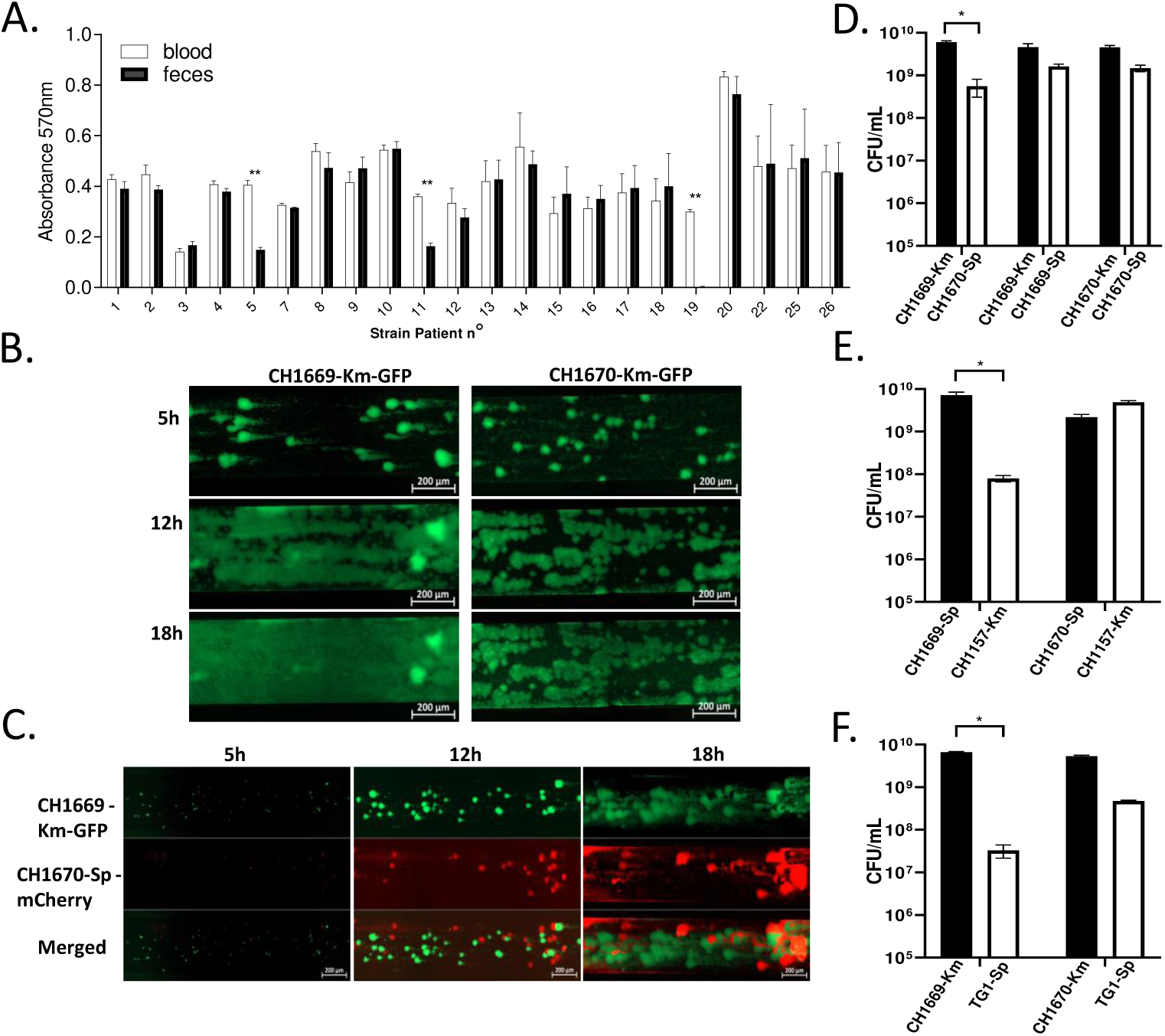
Distinct biofilm dynamics and improved bacterial competitiveness in an in vivo–adapted strain compared to a stool isolate. **(A.)** Biofilm biomass produced after 4h of culture in M63B1 medium by the paired strains of *Klebsiella* isolated from rectal carriage (stool) or bacteriemia (blood), stained by crystal violet (Absorbance 570nm). Specific biofilm structure for concordant paired strains isolated from patient #5, CH1669 isolated from blood sample and CH1670 isolated from rectal swabbing alone **(B.)** and co-cultured **(C.)** Strains were followed in the BioFlux microfluidic system at 37°C under a shear force of 0.5 dyn/cm² using epifluorescence microscope (Axio observer 7, Zeiss) at a magnification of 20×. Images were acquired in real time at T = 5 h, 12 and 18h. CH1669, isolated from blood, competitiveness against CH1670 isolated from rectal swabbing **(D.)**; *K. pneumoniae* CH1157 **(E**.) and *E. coli* TG1 **(F.)**. Each value is the mean ± SEM of at least three separate experiments. Statistical analysis: nonparametric Mann-Whitney test between paired strains: *p < 0.05, **p < 0.01.

The significant differences observed in patient #5 were further confirmed by microfluidic kinetic analysis sine the strain isolated from the patient’s blood culture (CH1669-Km-GFP) produced more biofilm than the strain isolated from stool**s** (CH1670-Km-GFP). In addition, the blood culture isolate showed more homogeneous surface colonization with fewer aggregates, suggesting a distinct mode of biofilm organization (**Fig 5B**).

We then tested whether these changes could be adaptive and result in increased fitness. The same strains were placed in direct competition for biofilm formation in the microfluidic channel. Under these conditions, the blood culture isolate (CH1669-Km-GFP) progressively dominated the system after 18 h of growth compared to CH1670-Sp-mCherry. This competitive advantage, quantified by CFU counts, was significant at the inter-clonal (CH1669 versus CH1670) (**Fig 5D**), intra-species (CH1669 versus *K. pneumoniae* CH1477) (**Fig 5E**), and inter-species (CH1669 versus *E. coli* CH1786) (**Fig 5F**) levels. In contrast, the stool isolate CH1670 was not able to achieve the same level of competitiveness.

## Discussion

In this study, we traced *Klebsiella* from the gut to the bloodstream in ICU patients with bacteremia, to uncover within-patient evolutionary events that could link intestinal colonization to invasive infection. Understanding the association between asymptomatic *Klebsiella* colonization and subsequent disseminated infection could provide opportunities for identifying high-risk patients, and intervention and ultimately prevention of infection (8). In the cohort of patients analyzed in this study, exposure to antibiotics within 2 weeks prior to bacteremia (95.5% of patients) was the most relevant risk factor of infection, as previously observed in the literature (27). The risk of developing a sepsis within 90 days when patients are exposed to high-risk antibiotics (3rd/4th-generation cephalosporins, lincosamides, fluoroquinolones, beta-lactam/beta-lactamase inhibitor combinations, vancomycin and carbapenems) was 65% higher than in those without antibiotic exposure (28). All different antibiotics and medications administered in the ICU (such as proton pump inhibitors, for example) are likely to induce a dysbiosis of the gut microbiota and therefore the proliferation of antibiotic resistant *Klebsiella* within the intestinal microbiota. Members of the Proteobacteria, such as *E. coli* or *K. pneumoniae*, normally represent less than 2% of the microbiota, but these bacteria can account for up to 30% of the total species in a dysbiotic state (9). In the study performed by Shimasaki *et al*., high CR-KP load values were detected in patient rectal swab samples in the week before the onset of the bloodstream infection episode (10). This suggests either increased prevalence of KP strains or increased competitive abilities. Results obtained with patient #5 would indicate that the two events can occur simultaneously. We showed that acquisition of a plasmid results in increased survival upon antibiotic stress and increased competitive abilities of the blood sample compared to those of the fecal sample.

Of the five patients whose strains were sequenced, patients #1, #2, #4, and #5 had been exposed to antibiotics within two weeks prior to the onset of bacteremia. Patients #2, #4, and #5 specifically received beta-lactams during this period. This antibiotic exposure could have facilitated the acquisition of plasmid and chromosomal broad-spectrum beta-lactamase encoding gene *bla*SHV-1 in the strain isolated from the blood of patient**s** #4 and #5 compared to their fecal counterparts, and this acquisition could explain the increase in ampicillin resistance of the blood strains. Conversely, the absence of exposure to beta-lactam in the two weeks preceding the bacteriemia in patient #1 could have promoted the plasmid rearrangement that led to the loss of the *bla*OXA-1 copies in the strain isolated from blood compared to its fecal counterpart strain. Taken together, our findings confirm the rapid turnover of antibiotic resistance genes within patients, underscoring the remarkable genomic plasticity of *Klebsiella*, which can swiftly acquire and lose genes that provide transient advantages and subsequently discard them once they become a burden.

The evolution of *K. pneumoniae* coupled with MDR gene acquisition within a host continuously exposed to multiple antibiotics has been documented elsewhere (29, 30). In our study, analyses with Kleborate, the pipeline based on several databases of antibiotic resistance genes, detected the presence of resistance genes (*bla*, *oxq* or *qnr* family) in bloodstream isolates compared to the stool samples, either chromosome or plasmid. Increased AT richness and gene gain observed in blood isolates compared to their fecal respective isolates are strongly suggestive of horizontal gene transfer (HGT) events. Indeed, the presence of additional plasmids was observed in two blood isolates (CH1669 and CH1668) compared to their respective stool isolates. We hypothesized that acquisition of these plasmids was due to horizontal gene transfer events but our attempts to generate *in vitro* transfer by mating of plasmids from CH1669 strain to several recipient strain**s** (CH1670, *K. pneumoniae* Ch1157 or *E. coli* TG1) did not yield positive results (data not shown). The identification of a phage within the plasmid of strain CH1669 has been suggested as illustrating the contribution of virome-microbiota interactions to horizontal gene transfer, including the dissemination of ARGs (31, 32), but we detected no conjugation or canonical mobility genes on CH1669 plasmids. These observations suggest that the element mediates the transfer through plasmid-encoded mechanisms and is mobilized, at least partly, via its phage functions. In addition, the CH1668 8.8kb plasmid had a very high identity (98%) to the well characterized pSEM from *Salmonella enterica* serovar Typhimurium plasmid (25), which has been found in a wide range of enterobacteria and is mostly localized in the chromosome in the form of an ICE. The backbone of this ICE, present in the plasmids of all the other strains in this study, can therefore adopt different mobilizable strategies (ICE, plasmid, phage-mediated), suggesting its great plasticity and its potential for function exchanges, as suggested by the presence of sequences carrying resistance to silver, arsenic, and copper. A recent study showed that plasmids belonging to the *IncF* family mediate the clonal (within the same lineage) and horizontal (between lineages) transmission of the ESBL gene *bla*_CTX-M-15_ within *Klebsiella* strains isolated from blood samples (33). Further work is needed to unravel genetic strain adaptations and their impact on virulence. The mechanism of in situ horizontal gene transfer is particularly efficient for harmful bacterial pathogens, such as *Klebsiella*, which can acquire new resistance genes from co-existing commensals within the patient’s gut microbiota (34). However, *in vivo* development of antibiotic resistance may also be attributed to initial colonization of the patients by multiple strains rather than gene acquisition by one clonal strain, as shown by Ding *et al.* with carbapenem resistance Enterobacteriaceae (35).

In a previous study, we showed that long-term intestinal colonization and associated dysbiosis were linked to the presence of a type 6 secretion system (T6SS) in the pathobiont (36). A complete cluster encoding T6SS was present in the sequence genomes of all *Klebsiella* isolates in this study, and *Klebsiella* isolates from patient rectal swabs were able to adhere to intestinal epithelial cells. This close interaction was not associated with an invasion process of *Klebsiella*. To note, a conventional aminoglycoside protection test is not suitable to assess the invasion capacity of biofilm-producing bacteria, owing to the formation of aggregates that prevent the action of the antibiotic molecules. Additionally, none of the fecal isolates from this previous study was able to destabilize *in vitro* the trans-epithelial barrier, suggesting that the passage into the bloodstream is not associated with active trans- or paracellular passage. In a study describing an invasive *K. pneumoniae* strain isolated from a patient with bacteremia, the authors showed that its paracellular translocation involves the cellular Rho-GTPases and the phosphoinositide 3-kinase (PI3K)/Akt pathway, leading to modulation of microtubules and rearrangement of the cytoskeleton of the intestinal epithelial cell (19). Also, the toxinogenic *K. oxytoca* AHC6 strain has been shown to increase colic permeability in a murine model of intestinal colonization by decreasing tight junction protein expression (claudins 5 and 8), independently of cytotoxin production (18). These two processes (transepithelial or passage through the intercellular junctions) seem to be strain-dependent and are not necessarily a general phenomenon.

In the present study, patient #3 had no documented antibiotic exposure before the onset of bacteremia, which was identified at the time of ICU admission. Antibiotic pressure and dysbiosis alone do not seem to fully account for the transition from intestinal colonization to bloodstream infection and it is likely that the immune system also plays a major role in controlling the dissemination of *Klebsiella* within the organism. The immunodeficient profile induced by sepsis in most patients could also explain the translocation process of *Klebsiella* from the intestine to the blood. The growth arrest-specific 6 (Gas6), which is secreted by intestinal macrophages, has been shown in a murine model to inhibit *Klebsiella* translocation from the gastrointestinal tract by strengthening tight junction barriers in the intestinal epithelium (37). Further studies are required to elucidate the dysfunctional mechanisms of the immune system that contribute to increased susceptibility to *Klebsiella* infection, but it is also necessary to elucidate the bacterial changes induced within the intestinal environment that could lead to the selection of individual cells particularly capable of passing through the epithelium barrier.

In this study, we analyzed the whole genome sequences of five paired strains all phylogenetically related to one another in patients of whom none were infected with the same strain. SNPs were observed in all isolates from blood samples comparatively to their fecal counterpart, with a very high rate in strain CH1661 from patient #1 (323 SNPs detected). The acceleration of the genome evolution of this strain, probably caused by the mutation in *mutS*, refers to an hypermutator phenotype (38). The genome of this isolate also had mutations in genes encoding efflux pumps (AcrB) and resistance to several antimicrobials such as fusaric acid and beta-lactamase. Most single nucleotide changes observed in the genomes of bloodstream isolates occurred in genes associated either with surface structures and properties or with resistance to antibiotics. Such modifications have previously been associated with altered capsule export (52) and increased hypermucoviscosity (21, 39). The occurrence of SNPs in *wzc* has also been previously associated with increased hypermucoviscosity, a proxy of hypervirulence. For instance, the capsule of *K. pneumoniae* weakens the host’s immune response, particularly the maturation of dendritic cells (DCs) and the production of pro-Th1 cytokines (40), leading to better immune system escape. In contrast, in a study of the genomic dynamics of four ST11-CRKP sequentially isolated from urine samples, Ye *et al.* suggested that a reduction in polysaccharidic capsule in *Klebsiella* is associated with an adaptive evolution of virulence (41). Two types of capsule mutants, each associated with a specific pathology, have emerged: hypercapsulated mutants able to resist to phagocytosis, associated with bloodstream infections, and capsule-deficient mutants associated with urinary tract infections (42). Interestingly, KPC-Kp isolates in human serum undergo *wcaJ* loss-of-function mutation, which confers resistance to complement-mediated serum killing but also increased complement C3 binding and susceptibility to opsonophagocytosis (43). Together, these observations emphasize the complex and context-dependent role of capsule modulation in *K. pneumoniae* pathoadaptation, showing that both increased and decreased capsule expression can confer selective advantages in distinct host environments. Modifications of the capsule biophysical properties and/or altered expression of surface adhesins could facilitate the in vivo survival of the pathogen. The genome of strain CH1669 had a mutation in *cpxA*, which encodes a repressor of type I and type III fimbriae encoding genes (44), and a reversion of the *fim* switch. A significant increase in biofilm formation was observed for the blood isolate of **this duo**? Which pair? compared to **its**? their rectal swab counterpart (**Fig 5A**), accompanied by a distinct biofilm structure (**Fig 5B**), suggesting that a gain of function had occurred in the strain, enhancing its adhesive capacity during the potential migration from the intestinal reservoir to the bloodstream. In addition, this in vivo–adapted strain has strong inter- and intra-species competitive abilities, which could enhance its capacity to dominate both the host microbiota and potential pathogenic niches. The increased fitness observed in this strain can be attributed to the phage in its plasmid that is seen in the blood strain, but not in the fecal isolate. Prophages, which are common in intestinal bacteria, confer fitness advantages to their hosts, facilitate bacterial adaptation to new niches, and play a significant role in their evolution (45, 46). The fact that the strain with increased fitness carries a phage within a plasmid suggests a possible link with lysogenic phage induction. However, it has been shown that the presence of a plasmid containing conserved phage genes is correlated with a virulence reduction in *K. pneumoniae* (47). To further elucidate the strain’s virulence and the functional potential of its phage-derived mobile genetic element, additional studies will be required to characterize its transcriptional and translational regulation and to assess the virulence phenotype in appropriate infection models.

In conclusion, this work identifies possible within-host evolutions during multiple antibiotic treatment of a major nosocomial pathogen, mainly driven by HGT and few SNP on gene encoding surface elements linked to functional phenotypes of bacterial colonization and fitness. As the isolates were initially selected for their ability to resist antibiotics, future research needs to be carried out on initially susceptible strains and extended to different samples from disseminated infections in the patient (due to bronchoalveolar lavage, or urine or medical devices) to decipher which genetic determinants affect the eco-evolutionary success of clonal groups. A better understanding of the evolutionary capacity of bacteria is important for predicting the emergence of antimicrobial resistance within patients and for developing strategies for pathogen surveillance and infection control. An original strategy based on in situ editing of the bacterial genes within the gut microbiota has recently been developed in a murine model (48) and could pave the way for future microbiome-targeted therapies to prevent antibiotic failure.

## Material and Methods

### Bacterial strains and culture conditions

The *Klebsiella spp* strains used in this study, *K. pneumoniae* and *K. oxytoca*, were isolated between January 2017 and March 2021 from patients who had developed bacteremia in intensive care units. They were isolated as part of routine care, and no additional samples were taken as part of this study. Patients under the age of 18, under guardianship, curatorship, or who were pregnant were excluded. The study was approved by local Ethics Committee (IRB00013412, “CHU de Clermont Ferrand IRB #1”, IRB number 2022-CF017) and complied with the French policy of individual data protection. For each patient, *Klebsiella spp*. collected from rectal swabs and blood samples were retained when they had similar matrix-assisted laser desorption ionization time-of-flight (MALDI TOF) and enterobacterial repetitive intergenic consensus (ERIC)-PCR amplification profiles (49). Strains were preserved by freezing at -80°C in 20% glycerol, and then cultured for manipulations at 37°C with agitation in lysogeny broth (LB) medium. The equivalence between colony-forming units (CFU) and absorbance at 620nm (A_620nm_) was determined for each strain.

To construct fluorescent strains, GFP or mCherry encoding genes were introduced into the chromosome of the clinical *K. pneumoniae* strains using a mobilizable mini-Tn7 base vector by triparental mating involving *E. coli* MFD*pir* harboring the plasmid pUC18R6KT-mini-Tn7T-Km-*gfpmut3*, pUC18R6KT-mini-Tn7T-Sp-*mCherr*y *E. coli* MFDpir(pTNS3) and the recipient strain (50). The following derivate strains were obtained: CH1669mini-Tn7T-Km-*gfpmut3* (CH1669-Km-GFP), CH1669mini-Tn7T-Sp-*mCherr*y (CH1669-Sp-mCherry), CH1770mini-Tn7T-Km-*gfpmut3* (CH1670-Km-GFP) and CH1670mini-Tn7T-Sp-*mCherr*y (CH1670-Sp-mCherry).

For bacterial competitiveness assays, bacterial cultures were adjusted to an absorbance at 620 nm (A₆₂₀) of 0.6. Three volumes of a strain for one volume of another strain were mixed and spotted onto a sterile nitrocellulose filter placed on LB agar plates. After 24 h of incubation at 37 °C, the bacterial mixture was recovered from the filter, resuspended in sterile saline, and plated onto selective LB agar containing either kanamycin (Km), spectinomycin (Spec), or ampicillin (Amp) to determine the number of each strain. The strains used for competition were *K. pneumoniae* CH1669-Km-GFP (CH1669 mini-Tn7T-Km-*gfpmut3*) (51) or *E. coli* TG1-Sp (TG1mini-Tn7T-Sp-*mCherr*y). Competitiveness was assessed by calculating colony-forming units (CFU) recovered on selective media.

### Cellular cell lines and infection condition

The epithelial colon human adenocarcinoma cell line Caco-2 was cultured in Dulbecco’s modified Eagle medium (DMEM) with 10% decomplemented fetal calf serum (FCS) for 30 minutes at 56°C. All cell cultures were incubated at 37°C with 5% carbon dioxide in the presence of antibiotics (spectinomycin 100μg/ml and penicillin 100 IU/mL) until infection.

For adhesion experiments, Caco-2 cells were seeded in 24-well plates at 1×10^6^ cells per well for 24h. Prior to infection, the medium was changed to DMEM with 10% decomplemented FCS and no antibiotics. Infections were carried out with 1×10^7^ bacteria per well, giving a multiplicity of infection (MOI) of 10. The wells were then incubated at 37°C and 5% static CO_2_ for 4h. After this incubation time, the medium was removed and the wells were washed twice in phosphate-buffered saline (PBS). Cells were lysed with 1% Triton solution, and the number of adherent bacteria were quantified by plating serial dilutions on agar plates.

*Klebsiella spp* invasion capacities were assessed using Caco-2 epithelial cells: the infection step described above was followed by a 4h adhesion step, cells were washed twice with PBS and incubated for 1h at 37°C 5% static CO_2_ in DMEM culture medium with 10% FCS supplemented with amikacin (1000μg/mL). After two further PBS washes, the cells were lysed with a 1% Triton solution and the viable bacteria quantified by plating the suspension on agar plates.

### Biofilm formation

Bacteria were precultured for 18h and seeded (10^6^ CFU/mL) in 96-well flat-bottom plates in 100μl of M63B1 minimum medium with 0.4% (v/v) glucose, and incubated for 4h in a static oven at 37°C. The medium was then removed and the biofilms formed at the bottom of the wells washed with 100μL of PBS. Staining with 0.5% crystal violet was performed for 10 minutes at room temperature (Magana *et al.*, 2018). The dye was dissolved with 200μl of ethanol per well, and biofilm biomass was measured with a spectrophotometer at 570nm (Epoch® Reader Microplate Spectrophotometer). A second test was performed to estimate biofilm resistance to amikacin as follows; biofilms were formed for 4 h in 24-well plates after seeding with 1×10^7^ bacteria per well, in DMEM medium with 10% FCS decomplemented without antibiotics. After washing the medium, amikacin 1mg/ml was added for 1 hour. Live bacteria were then counted as above.

Biofilm development was studied using the BioFlux™ 200 microfluidic system (Fluxion Biosciences, South San Francisco, CA, USA) with 48-well plates as previously described (51). Channels (75 μm depth, 350 μm width) were primed for 5 min with 500 μL of M63B1-0.4% Glc at 5 dyn/cm². *K. pneumoniae* overnight cultures in M63B1-0.4% Glc were adjusted to A₆₂₀_nm_ 0.01 (∼10⁷ CFU/mL) and inoculated by reverse flow at 5 dyn/cm² for 2 seconds. After a 15 min adhesion step at 37 °C without flow, biofilm growth was monitored at 37 °C under a shear force of 0.5 dyn/cm² (63 μL/h) using an Axio Observer 7 inverted epifluorescence microscope (Zeiss, 20×). Images were acquired every hour for 18 h and analyzed with Zen 2 software (Zeiss). Biofilm mechanical strength was tested after 18 h by increasing shear force from 0.5 to 10 dyn/cm².

### Measurement of transepithelial electrical resistance (TEER)

Transwells (Merck® Millipore) with 0.4-μm pore size and 0.33-cm^2^ filtration area were seeded with 1×10^6^ Caco-2 epithelial cells in 12-well plates. Four hundred µL of DMEM medium with 10% FCS was added to the Transwell and 1,200μl to the well supporting it. Culture was performed in a 37°C 5% CO_2_ static oven, with the culture medium being changed every 48 hours for at least 14 days to obtain sufficiently differentiated epithelial cells. The culture was extended as long as the measured transepithelial electrical resistances were less than 600 Ohms per well (52). Resistances were measured with a Millicell ERS-2 V-Ohm meter (Merck® Millipore). Once resistance was reached, the wells were infected with 1×10^7^ bacteria per well (MOI 10). Resistances were measured every hour up to 6 hours after the start of infection. A positive control treatment was performed with 25μM H_2_O_2_ (53).

### Antimicrobial Susceptibility Test

Various breakpoint antibiotic resistances were determined by VITEK 2 Biomerieux liquid antibiotic susceptibility test: AM: Ampicillin; AMC: Amoxicillin/Clavulanic acid; TIC: Ticarcillin; TZP: piperacillin/tazobactam; FOX: Cefoxitin; CTX: cefotaxime; CAZ: ceftazidime; ETP: ertapenem; IPM: imipenem; TM: tobramycin; AN: amikacin; GM: gentamicin; NA: nalidixic acid; OF: ofloxacin; CIP: ciprofloxacin; SXT: trimethoprim sulfamethoxazole; FT: furanes (54). The CMI of ampicillin was determined using the broth micro-dilution method as adapted by the European Committee on Antimicrobial Susceptibility Testing (EUCAST) (54).

### Whole-genome sequencing and sequence analyses

***i) Genome preparation*** Genomic DNA (gDNA) was extracted from 45mg of bacterial pellet cultivated overnight in LB using the NucleoSpin® Microbial DNA kit (MACHEREY-NAGEL) following the manufacturer’s instructions. Complete long-read sequencing and assembly of gDNA from individual strains were performed with the Eurofins Genomics service using Oxford Nanopore Technologies (ONT). ***ii) Genome circularization***. Raw nanopore sequencing reads were assessed for quality and filtering. Filtlong v0.2.1 was used to remove short and low-quality reads. The assembly was performed using Flye v2.9.3 with parameters optimized for bacterial genomes. The resulting contigs were further polished using Medaka v1.8 to improve base accuracy. The quality of the assembled genomes was assessed with various quality assessment tools (QUAST v5.2, CheckM2 v1.0.1, Mash v2.3). Raw data were deposited in the SRA (Sequence Read Archive) database under the accession number PRJNA1159795. ***iii) Variant analyses***. Each bloodstream sample was compared to its respective stool sample sequence using the -cntg option of Snippy (55). For consistency, the blood samples were also compared with their respective fecal samples. To verify these results and further analyze genome rearrangements, large deletions, and unmapped sequences we also used *breseq* (0.30.1) with default parameters (56). ***iv) Klebsiella typing.*** Kleborate (v2) (57) was used to determine sequence type and K- and O-loci, and to characterize the different isolates in terms of virulence factors. To detect and annotate antibiotic resistance genes (ARGs), we used ResFinder v4.6 (58). Macsyfinder v2 (59) and the package T6SS were used to identify the presence of Type 6 secretion systems (60). ***v) Phylogenetic tree.*** To build a phylogenetic tree, we first identified the pangenome (*i.e.* the full repertoire of homologous gene families) using the module pangenome of the software PanACoTa v1.3.1 (61). Briefly, gene families were built with MMseqs2 with an identity and bidirectional coverage threshold of 80%. This analysis identified 8169 gene families We then computed the strict core genome, meaning that a gene family must be present in single copy in all genomes to be considered core, and found 3893 gene families. To compute the phylogenetic tree, we aligned each of the 3893 protein families of the persistent genome individually with the align module of PanACoTA. These alignments were concatenated to produce a large alignment matrix with 54737 parsimony-informative sites over a total alignment of 3,738,906 bp. We then used this alignment to make the phylogenetic inference using IQ-TREE multicore version 2.3.0 (62). The option -m TEST to search for the best likelihood method was used. The final tree was built using a GTR+F+I model, chosen according to Bayesian Information Criterion and calculated 1000 ultra-fast bootstrap. We rooted the phylogenetic tree with the midpoint root function from the *ape* R package. ***vi) MGE characterization*.** Plasmids were annotated with Bakta (63). To compare with NCBI genomes, the whole sequence was blasted with -max_target_sequences set to 5000.The distribution of the alignment lengths was analyzed. For plasmid in CH1668, those with less than 5000 bp aligned were filtered out. Plasmid visualization was performed using Prokasee (64). Additionally, Phigaroo, CARD and pLAnnotate included in Prokasee were also used to highlight plasmid features of interest. Phage sequences were confirmed by PHASTEST (65). Genetic comparison between mobile genetic elements was performed using clinker (66).

### Statistical tests

Statistical analyses were made with GraphPad Prism® software or R v4.3.1. The non-parametric Mann-Whitney or Kruskal-Wallis tests, with Dunn’s correction for multiple comparisons, were used. A p-value of less than 0.05 was considered significant.

## Acknowledgements

We thank the Auvergne Rhone Alpes Regional Health Agency for awarding a research grant to fund Cécile Gosset’s participation in this project. We are grateful to the Culturomic UCA Partner platform for its assistance and the resources provided. We also thank Manon Caudal and Hugo Rey-Dépreux for their technical help during their training course.

## Legends

**Figure S1: Representative ERIC-PCR fingerprint of different *Klebsiella* isolates on 1% agarose gel. (A.)** Discordant isolates. **(B.)** Concordant paired strains isolated from patient #5, CH1669 isolated from blood sample and CH1670 isolated from rectal swabbing. Primers used for ERIC-PCR amplification: ERIC 1 (5’-ATG TAA GCT CCT GGG GAT TCA-3’) et ERIC 2 (5’-AAG TAA GTG ACT GGG GTG AGC G-3’) (49).

**Figure S2: Characteristics of plasmids from strain CH1661 isolated from blood sample of patient** #**1. (A.)** and **(B.)** Plasmid annotation and visualization of strain CH1661. Plasmids were annotated with Bakta, and visualized with Prokasee. Additionally, Phigaroo, CARD and pLAnnotate, included in Prokasee; were used to indicate plasmid features of interest. **(C.)** Plasmid alignment of CH1661 and CH1662 strains, isolated from blood sample and rectal swab from patient #1, respectively.

**Figure S3: Plasmid annotation and visualization of strain CH1663 isolated from blood sample of patient** #**2.** Plasmid was annotated with Bakta, and visualized with Prokasee. Additionally, Phigaroo, CARD and pLAnnotate, included in Prokasee, were used to indicate plasmid features of interest.

**Figure S4: Plasmid annotation and visualization of strain CH1665 isolated from blood sample of patient** #**3.** Plasmid was annotated with Bakta, and visualized with Prokasee. Additionally, Phigaroo, CARD and pLAnnotate, included in Prokasee, were used to indicate plasmid features of interest.

**Figure S5. Length of alignment of CH1668 plasmid (A) and CH1669 plasmid #2 (B) against NCBI sequences.** Most hits displayed very low coverage of the different plasmids. Vertical dashed line corresponds to the cut-off used for further analyses.

